# Fully Automatic Cell Segmentation with Fourier Descriptors

**DOI:** 10.1101/2021.12.17.472408

**Authors:** Dominik Hirling, Peter Horvath

## Abstract

Cell segmentation is a fundamental problem in biology for which convolutional neural networks yield the best results nowadays. In this paper, we present FourierDist, a network, which is a modification of the popular StarDist and SplineDist architectures. While StarDist and SplineDist describe an object by the lengths of equiangular rays and control points respectively, our network utilizes Fourier descriptors, predicting a coefficient vector for every pixel on the image, which implicitly define the resulting segmentation. We evaluate our model on three different datasets, and show that Fourier descriptors can achieve a high level of accuracy with a small number of coefficients. FourierDist is also capable of accurately segmenting objects that are not star-shaped, a case where StarDist performs suboptimally according to our experiments.

## Introduction

Imaging, identifying, and morpho-measuring cells and their subcellular compartments are often the first step of fundamental cell biology research and drug discovery^1^. Single cell segmentation is an important part of this, as it enables researchers to analyze the number of cells, their shapes, phenotypes or physiological state on microscopic images. In the recent years, artificial neural networks emerged as an effective tool for tackling segmentation tasks2, beginning with R-CNN^3^, followed by U-Net4 and Mask-R-CNN^5^, just to mention some of them. Recently, StarDist6 became a popular solution due to its better cell representation with star-convex polygons, instead of using per-pixel classification, or bounding boxes. One drawback of this approach is that the objects to be segmented have to be star-shaped and the rays used to represent them have to be equiangular, which can also lead to poor representation when working with just a few of them. To overcome this issue, Mandal et al. proposed SplineDist^7^, a modification of StarDist, which represents cells with spline contours that are characterized by control points. The network predicts these points without any restriction to their location. SplineDist can also tackle cases where the objects are not star-convex. In this paper, we present a neural network called FourierDist: also using a U-Net backbone and fully-connected layers as the aforementioned networks, our method predicts a vector of Fourier descriptors^8^, which in turn can be reconstructed into a contour. Fourier descriptors have a longstanding history of effectively describing shapes, dating back to the works of Zahn et al.9 Due to the relatively simple elliptical shapes that cell nuclei have, we can closely estimate them by using only a few descriptors. This makes FourierDist able to work effectively with a very small number of predicted coefficients, where SplineDist doesn’t give any solution and StarDist yields results of poor quality. Our evaluations show that besides this advantage, our method is competitive even when we increase the number of parameters, with StarDist achieving the highest scores as the number of parameters increase substantially. If the target objects are non star-convex, FourierDist outperforms both StarDist and SplineDist.

## Methods

### StarDist and SplineDist

StarDist is a segmentation method based on convolutional neural networks. It takes an input image, and predicts a star-convex polygon with *N* vertices for every pixel that are equiangular to each other. Besides that, it also predicts a probability value *p_ij_* for every pixel (*i, j*) so that pixels with low object probability can be ignored. Once a prediction has been made for every pixel, a non-maximum suppression (NMS) step ensures that overlapping candidates are filtered out. StarDist uses a standard U-Net backbone with an additional 3 × 3 convolutional layer with 128 channels and ReLU activation function. For the object probability prediction, a single-channel convolutional layer is used with a sigmoid activation function. The output layer that predicts *N* polygon vertex distances is of size *N* without any activation function. SplineDist modifies this architecture by representing cells as B-spline contours instead of polygons. A spline contour of order *n* is a piecewise polynomial function with a degree of *n* – 1. B-splines of a given order can be thought of as basis functions in the space of spline functions, which means that any piecewise polynomial function with a degree of *n* – 1 can be described by a linear combination of B-splines of order *n*. The “weights” of the B-splines are the control points, which describe the spline curve itself implicitly. In the SplineDist model, cubic B-splines are used (meaning *n* = 4), and a cell is described by a function

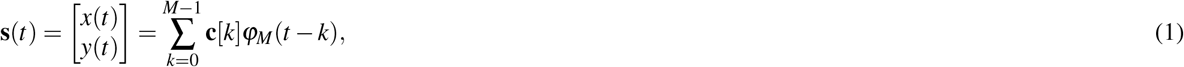

where *M* is the number of control points used, **c** is the vector of control points, and 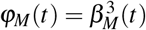 are the cubic basis functions for M number of control points. The SplineDist architecture only predicts the control points for every pixel (*i, j*) (besides the object probabilities *p_ij_*) that don’t have any restriction in terms of relative position to each other. This representation is beneficial because with a small number of parameters, smooth contours can be predicted, which is not the case for StarDist. One key difference from StarDist is that instead of calculating the distances of the predicted control points from the “ground truth” control points (which wouldn’t be possible, since the same contour can be expressed with a different set of control points and basis functions) in the loss function, a continuous contour is generated from the predicted {*c_ij_* [*k*]}_*k*=0,…,*M*–1_ points, which is then compared to the ground truth segmentation.

### Fourier descriptors

Using a set of basis functions in the space of contours and then taking the linear combination of them to describe shapes has been studied for a long time^9^. Besides the aforementioned basis spline functions, complex exponential bases and Fourier coefficients serve a similar purpose. Assume 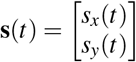 is a clockwise-oriented simple closed curve, where *t* ∈ [0,1]. In this case, **s** can also be written as a complex-valued function (*s*(*t*) = *x*(*t*) + *iy*(*t*)). Fourier’s theorem states that any contour such as s can be expressed with the following infinite series:

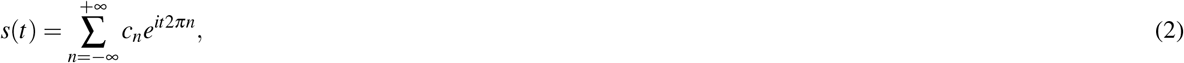

where *e*^*it*2*πn*^ are the basis functions describing circular motion alongside a unit circle with frequency *n*, and 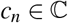 are the so-called “Fourier coefficients”, that control the radius of the circle and the starting position of the n-th circulation expressed by the exponential. The exponential terms are fixed for every contour, so the only parameters that can be changed in order to construct a different curve are the *cn* coefficients. Compared to the B-spline representation, where the same curve can be represented by multiple number of basis functions and control points, the Fourier basis functions form an orthogonal basis in the space of contours, meaning that any contour can be explicitly expressed with a linear combination of them. We can calculate all of the Fourier coefficients (the weights for the basis functions) explicitly with the formula

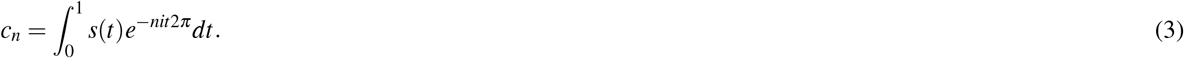

This description would not be useful if we needed all of the basis functions to properly describe a contour, since the number of coefficients needed for that would be infinite. However, by just using a few terms located around the 0-th exponential term, we can get a close approximation of the curve to be described. By utilizing this infinite series, an approximation for the curve s of order *N* is the following:

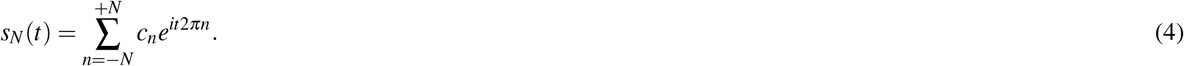

As the number of coefficients increases, the description becomes more accurate, however, terms further away from the 0-th exponential describe components of higher frequency and do not contain as much information as those of lower frequency. An example of the increasing reconstruction accuracy compared to StarDist can be seen on Figure 1.

**Figure 1.**
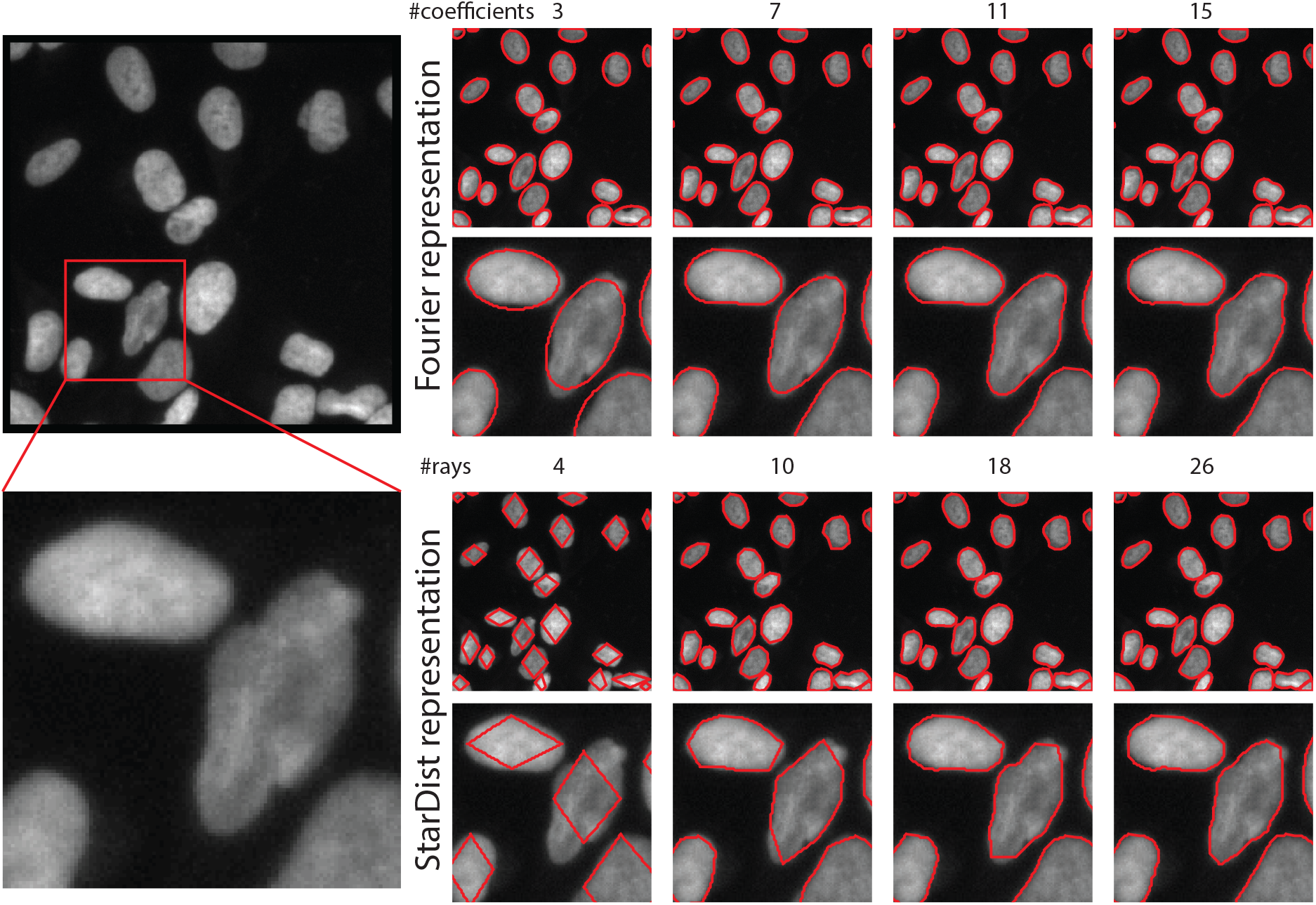
Reconstruction capabilities of the StarDist representation and Fourier descriptors. For just a few number of parameters, StarDist produces contours of low resolution. Fourier descriptors on the other hand already yield acceptable contours for the minimal number of parameters. Both representations increase in accuracy as the number of parameters increase.

### The FourierDist method

For every pixel (*i, j*) we regress the coefficients of order 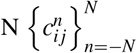 corresponding to the contour describing the cell, thus getting a prediction map 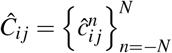. We make one slight modification on one of the predicted coefficients: instead of predicting 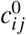 explicitly (which is the center of mass of the predicted curve), we train our model to learn 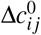, which describes the translation of the center of mass of the object relative to the pixel (*i, j*). Besides predicting a coefficient vector for every pixel, we also predict the probability of the pixel belonging to an object (cell) denoted by 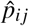. The proposed architecture can be seen on Figure 2. For training, we use the following loss function: for the object probabilities, we use the binary cross-entropy loss, as for the Fourier coefficients, we calculate the mean absolute error of the predicted and ground truth descriptors:

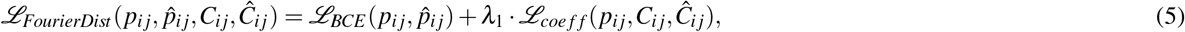

where

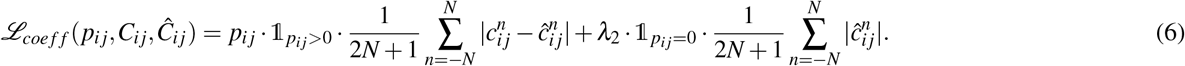

**Figure 2.**
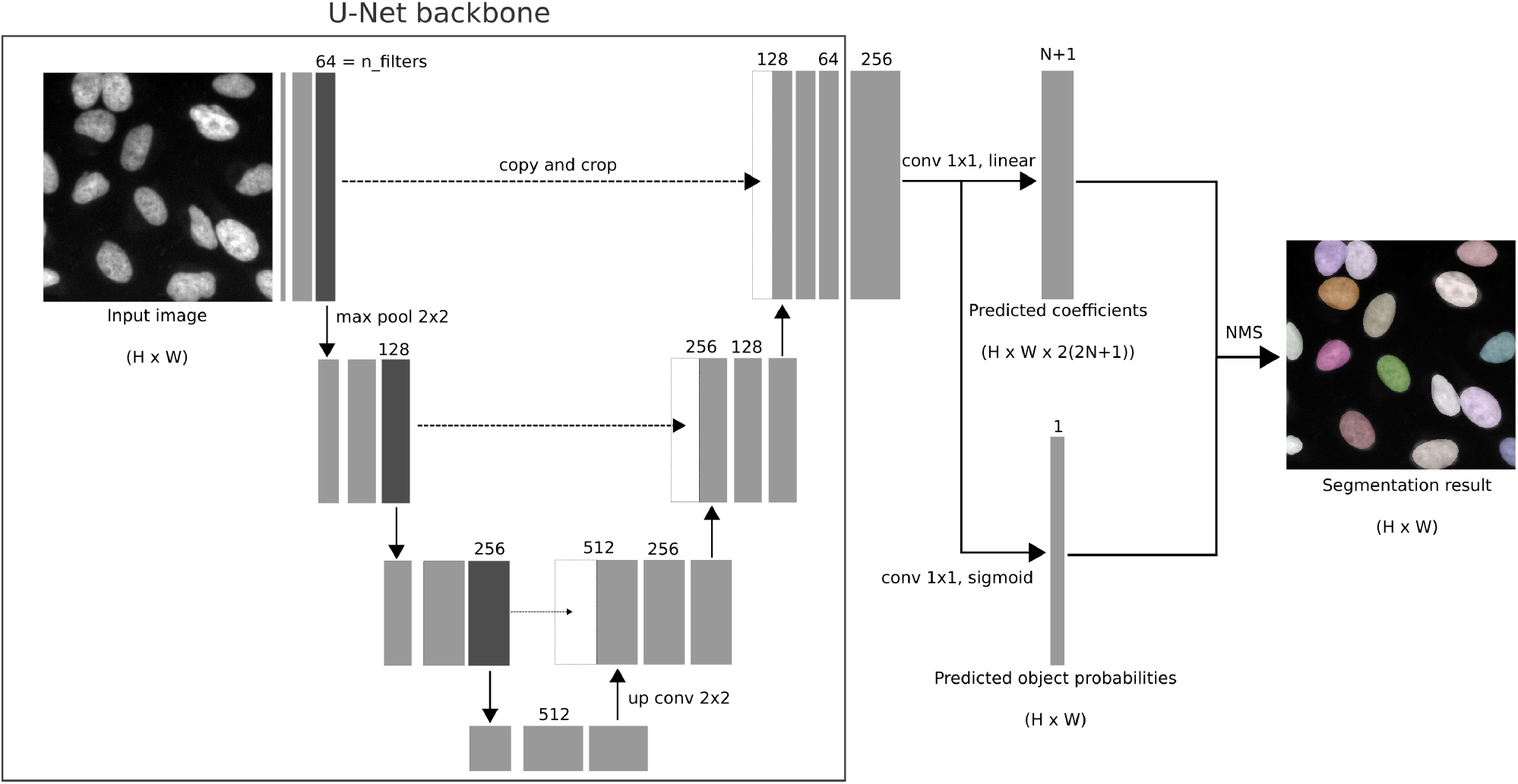
Architecture of the FourierDist network. We use a U-Net backbone with a depth of 3. After the last feature layer, we add a 3 ×3 convolutional layer, after which the object probabilities are calculated by a 1 × 1 convolutional layer with *sigmoid* activation. The coefficient vectors are calculated by a 1 × 1 conv layer with *linear* activation. Finally, the overlapping candidates are filtered out by a non-maximum suppression step.

## Results

For evaluating the efficiency of our model, and comparing it to other, state of the art methods, we used three different datasets: the DSB201810 (2018 Kaggle data science bowl dataset), a common benchmarking image set, the TCGA - The Cancer Genome Atlas histopathology dataset^11^, which consists of various tissue samples, and a synthetic (non star-convex) dataset, which was generated by the authors of SplineDist^7^. In order to make the results comparable to each other, we used the same hyperparameters and augmentation strategy for training the models (Adam optimizer with *λ* = 3 × 10^-4^, batch sizes of 4, and 300 epochs). For quantitative evaluation, we used the mAP (mean average precision) metric:

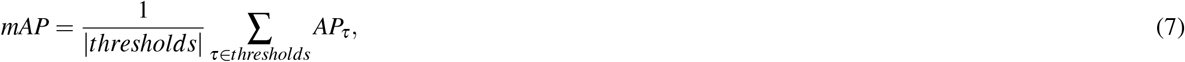

where *thresholds* = {0.5,..., 0.9}, and *AP_τ_* is the average precision for IoU threshold *τ*. The number of parameters for the networks are determined by the number of rays (R), two times the number of points (2 × *M*) and twice the number of coefficients (2 × (2*N* + 1)) for StarDist, SplineDist and FourierDist, respectively. For SplineDist, we use cubic splines (*β*^3^) as proposed by the authors. In Table 1, we quantitatively compare the efficiency of StarDist, SplineDist and FourierDist, with increasing number of parameters. As expected, StarDist performs poorly when the number of parameters are low (*R* ≤ 10), but gradually increasing. SplineDist and FourierDist on the other hand already start with higher scores, but the performance doesn’t increase substantially with increasing number of points/coefficients. An explanation for this is that for Fourier descriptors, numerically approximating the higher-order coefficients can lead to additional noise that can reduce the performance of our method. In the DSB2018 and TCGA datasets, FourierDist yielded the highest scores for smaller prediction vector sizes (6 and 10 parameters), while StarDist performed the best for 22 and 34 parameters. On the other hand, in the synthetic dataset, FourierDist produced the highest scores across all prediction vector sizes. Overall, we conclude that the advantage of FourierDist is that it can produce accurate segmentations even for very small prediction vector sizes. As the number of parameters increase, StarDist eventually will outperform both SplineDist and FourierDist. However, for a dataset that contains objects which are non star-convex, FourierDist will outperform StarDist in every scenario. We also note that StarDist requires the least amount of parameters in order to be able to function (*R* = 4) followed by FourierDist (2(2*N* + 1) = 6) and SplineDist (2*M* = 8). Examples from the three different datasets can be seen in Figure 3.

**Table 1.** Nucleus detection results for three different datasets for 4 different types of parameter settings. StarDist, SplineDist, and FourierDist are compared to each other.

| nparams | 6 | 10 | 22 | 34 |
| --- | --- | --- | --- | --- |
| DSB2018 |  |  |  |  |
| StarDist | 0.4359 | 0.5686 | **0.6037** | **0.6091** |
| SplineDist | - | 0.5662 | 0.5925 | 0.5689 |
| FourierDist (Proposed) | **0.5629** | **0.5745** | 0.5883 | 0.5784 |
| TCGA |  |  |  |  |
| StarDist | 0.2338 | 0.2985 | **0.3243** | **0.3413** |
| SplineDist | - | 0.2978 | 0.3070 | 0.2865 |
| FourierDist (Proposed) | **0.3142** | **0.3112** | 0.3145 | 0.2964 |
| Synthetic |  |  |  |  |
| StarDist | 0.3720 | 0.5390 | 0.6488 | 0.6654 |
| SplineDist | - | 0.5883 | 0.6392 | 0.6394 |
| FourierDist (Proposed) | **0.4591** | **0.6077** | **0.6998** | **0.6771** |

**Figure 3.**
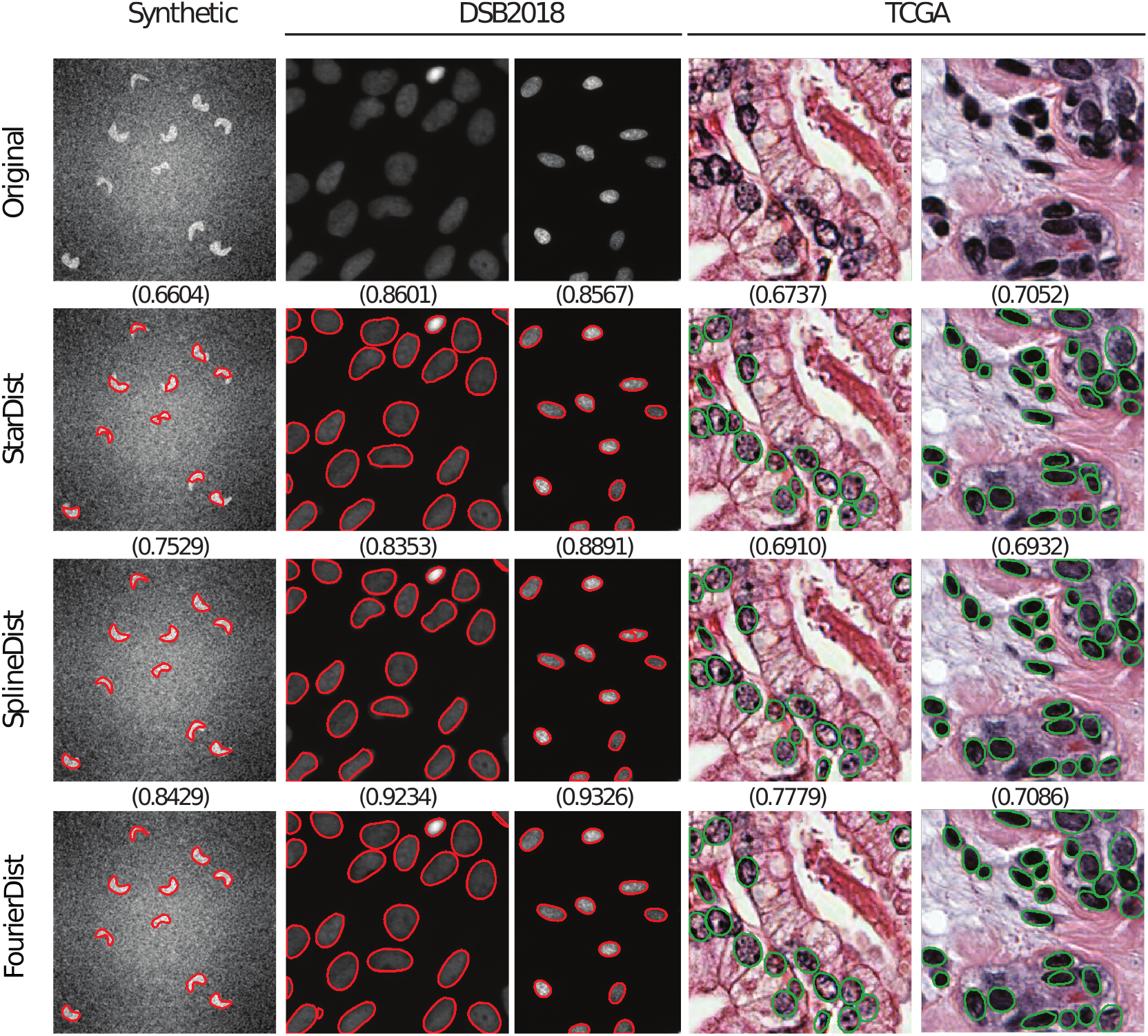
Sample segmentation results with equal number of parameters (*R* = 2*M* = 2(2*N* + 1) = 14) on the DSB2018, TCGA, and synthetic datasets. The AP metric is used for evaluation with an IoU threshold of 0.5, which can be seen above the images.

## Discussion

We proposed FourierDist, a neural network architecture based on StarDist, which predicts object probability values *p_ij_* and corresponding Fourier coefficients *c_ij_* for every pixel (*i, j*) on a target microscopy image. The lower-order Fourier descriptors can capture cell-shaped objects very well, hence the proposed model can achieve high segmentation accuracy for a very low number of parameters, a domain where StarDist does not perform well. Our quantitative results show that while clearly having an advantage with a low number of parameters, our model is slightly outperformed by StarDist for higher number of parameters. However, for synthetic examples, where objects are not star-convex, FourierDist clearly outperforms both StarDist and SplineDist for every examined prediction vector size. Our research opens up interesting possibilities for the future: studying other sets of basis functions that could be used to further enhance the segmentation accuracy in microscopy images, outperforming state-of-the art methods for every prediction vector size is something that is worth investigating.

## Acknowledgements

We acknowledge support from the LENDULET-BIOMAG Grant (2018-342), from the European Regional Development Funds (GINOP-2.3.2-15-2016-00006, GINOP-2.3.2-15-2016-00026, GINOP-2.3.2-15-2016-00037), from the H2020 (ERAPERMED-COMPASS, DiscovAIR), and from the Chan Zuckerberg Initiative (Deep Visual Proteomics). Prepared with the professional support of the Doctoral Student Scholarship Program of the Co-operative Doctoral Program of the Ministry of Innovation and Technology financed from the National Research, Development and Innovation Fund.

## Author contributions statement

D.H. designed the network architecture, implemented the method, and evaluated the results. P.H. supervised the project. All authors edited the figures and reviewed the manuscript.

## Additional information

The authors declare no competing interests.

Code availability: https://bitbucket.org/biomag/fourierdist

